# QCatch: A framework for quality control assessment and analysis of single-cell sequencing data

**DOI:** 10.1101/2025.06.15.659779

**Authors:** Yuan Gao, Dongze He, Rob Patro

**Author notes:** These authors contributed equally (Yuan Gao, Dongze He).

## Abstract

**Motivation:** Single-cell sequencing data analysis requires robust quality control (QC) to mitigate technical artifacts and ensure reliable downstream results. While tools like `alevin-fry` and `simpleaf` (and augmented execution context for the `alevin-fry`), offer flexibility and computational efficiency to process single-cell data, this ecosystem will further benefit from a standardized QC reporting tailored for its outputs.

**Results:** We introduce `QCatch`, a Python-based command-line tool that generates comprehensive and interactive HTML QC reports designed specifically for single-cell quantification results. Taking the output directory of `alevin-fry` or `simpleaf` as the input, `QCatch` is able to perform essential processing steps, like cell calling, and generate detailed QC reports that contain informative visualizations and statistics, including unique molecular identifier (UMI) count distributions, sequencing saturation estimates, and splicing status information, for QC assurance. Built for seamless integration into downstream analysis workflows, `QCatch` exports the processed results in a richly-annotated `H5AD` format file, a widely used data format common among many downstream single-cell data analysis tools.

**Availability:** The source code and documentation of `QCatch` are available on GitHub at https://github.com/COMBINE-lab/QCatch. `QCatch` can be installed via both Bioconda and PyPI.

## Introduction

Quality control (QC) of pre-processed data is a critical step in sequencing data analysis. It helps ensure data integrity by identifying technical artifacts, failures and biases introduced during sample preparation and library sequencing. This process is especially important in large-scale single-cell RNA-seq studies, where data quality directly influences the accuracy and robustness of downstream analyses.

As with most technologies, QC assessment in single-cell RNA sequencing (scRNA-seq) typically relies on detailed, comprehensive QC reports. While general-purpose QC tools exist [1], many scRNA-seq processing tools include their own reporting modules, enabling QC summaries that are tailored to their specific quantification outputs. For example, Cell Ranger [2] generates a web-based QC report that includes not only standard metrics but also highlights potential issues by displaying warnings when signs of low-quality or failed samples are detected for 10X Genomics assays. Recently, we developed a software ecosystem for single-cell data processing applicable for a wide range of UMI-based assays, accessible through the simplified command-line interface provided by `simpleaf` [3]. This ecosystem is built on top of `alevin-fry` [4], a fast, accurate, and memory-efficient tool for splice-aware quantification of single-cell sequencing data. By setting flags for splice-aware quantification, `alevin-fry` separately reports spliced and unspliced transcript counts for genes, making it suitable for a wide range of downstream analyses.

Despite the streamlined interface provided by `simpleaf` for end-to-end raw data processing, and the excellent performance of `alevin-fry`, and various backend tools such as piscem (https://github.com/COMBINE-lab/piscem), salmon [5] and cuttlefish [6], this ecosystem currently lacks a feature-rich post-processing module for QC reporting and other essential tasks such as cell calling, sometimes also called empty droplet detection. This gap is particularly notable when assessing splice-aware quantification results. Although AlevinQC (https://github.com/csoneson/alevinQC), which was designed for results produced by alevin [7], has been extended to support some outputs from `alevin-fry`, a more comprehensive and fully-integrated QC solution tailored specifically to `alevin-fry` is still needed.

To meet the growing demand from our user community, we have developed `QCatch`, an open-source, Python-based command-line tool specifically designed to support a range of post-processing tasks for quantification results generated by `alevin-fry`. As the newest addition to our single-cell software ecosystem, `QCatch` fills an important gap in our single-cell data analysis pipeline by performing essential processing steps including cell calling. Furthermore, `QCatch` provides a comprehensive set of QC metrics tailored for both traditional and splice-aware quantification by `alevin-fry`, presented through easy-to-read tables and interactive visualizations in the resulting QC report.

The open-source, modular architecture of `QCatch` promotes transparency, ensures reproducibility, and enables usability and adaptability across diverse single-cell sequencing datasets from a wide range of UMI-based assays. Prioritizing usability and comprehensiveness, `QCatch` produces visually intuitive, interactive QC reports that facilitate accurate evaluation of data quality. These reports empower researchers to interpret quantification results with greater confidence, supporting more robust biological conclusions. Designed with user-friendliness in mind, `QCatch` accepts the output directory of either `alevin-fry` or `simpleaf` as input and outputs both an HTML report and a richly-annotated H5AD object (one of the most widely used on-disk formats for single-cell data storage). This design choice enables seamless integration with the scVerse ecosystem [8], which currently includes over 80 tools for single-cell and spatial transcriptomics data analysis, such as scanpy [9].

## Methods

### Datasets

In this study, we applied `QCatch` to a publicly available dataset of human peripheral blood mononuclear cells (PBMCs) obtained from the 10X website at https://www.10xgenomics.com/datasets/1-k-pbm-cs-from-a-healthy-donor-v-3-chemistry-3-standard-3-0-0. This dataset comprises scRNA-seq data from 1k PBMCs isolated from a healthy donor, generated using the Chromium Single Cell 3’ Gene Expression v3 chemistry.

### Design and Implementation

`QCatch` is an open-source Python package available on GitHub (https://github.com/COMBINE-lab/QCatch), and can be installed via Bioconda [10] or PyPI [11]. Designed for command-line use, `QCatch` takes as input a `alevin-fry` or `simpleaf` output directory to generate a comprehensive QC report by automatically locating the quantification results and log files within that directory. Furthermore, `QCatch` exports cell calling results to a modified H5AD file or a user-specified output directory to facilitate downstream analysis.

### Functionality

`QCatch` first loads the quantification results from the input directory to assemble an AnnData object using the pyroe package version 0.9.0. The default count layer, X, contains the total UMI counts. For splice-aware quantification results generated by `alevin-fry`, in which spliced, unspliced and ambiguous UMI counts are reported separately, `QCatch` generates the X layer using the sum of all three types of UMI counts, and also stores them in different layers, namely the S, U and A layers, respectively. Then, `QCatch` parses the log information from the “quantification” and “generate permit list” steps of `alevin-fry`, using the respective log files. This information is summarized in a tabular form under the “Log info” section of the resulting QC report. If the input directory is generated by `simpleaf` version 0.19.3 or later, which exports the entire data package as a H5AD file, `QCatch` will directly load the H5AD file using the scanpy package (v1.10.4).

`QCatch` implements a customized cell calling algorithm based on the methodology established in CellRanger. This algorithm consists of two primary components: Order of Magnitude (OrdMag) estimation and EmptyDrops [12]. The OrdMag-based method, inspired by a thorough discussion with the EmptyDrops team (https://github.com/MarioniLab/DropletUtils/issues/88), provides an initial estimate of the set of high-quality, live cells present in the experiment, based on the distribution of UMI counts of observed barcodes. Barcodes identified by this method are considered reliable, and are directly included as the initial set of retained cells, bypassing the subsequent filtering performed by EmptyDrops.

Specifically, the OrdMag method estimates the expected number of high-quality cells based on the distribution of UMI counts across all observed barcodes. It samples a series of integer values, 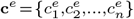, where each 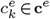 represents a candidate number of true cells. Given a full barcode list **b**, sorted in descending order by UMI count, for each 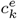, **b**_**k**_ represents the top 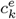 barcodes from **b**. From this subset, *m*_*k*_ is defined as the 99th percentile UMI count. This value is then used to define a dynamic cutoff for distinguishing true cells from background barcodes:

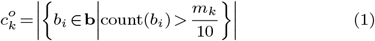

where count(*b*_*i*_) denotes the number of UMIs associated with barcode *b*_*i*_. In other words, 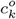 represents the observed number of rue cells corresponding to the candidate value 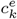 —that is, the number of barcodes in **b** with UMI counts greater than *m*_*k*_*/*10.

Thus, for each candidate 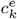, there is a corresponding observed count 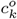, and the OrdMag method selects the expected number of cells by minimizing the loss function:

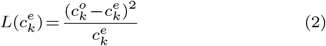

Subsequently, `QCatch` leverages a custom EmptyDrops implementation [13] to call additional cells from all cells except those in the primary set identified by OrdMag, and those having a UMI count less than 500. It calls cells by distinguishing true cells from the ambient RNA profiles using a multinomial model. The FDR threshold for calling a barcode as a cell is either 0.01 or 0.001, depending on the chemistry used. To note that, from the description, this method mimics the behavior of CellRanger version 9.0.

A basic qualitative visual representation is made available via UMAP [14] and tSNE [15] to project the high-dimensional gene expression data into a 2D space. To produce these visualizations, we adopt a standard pipeline, implemented using scanpy with default settings, which includes normalization, log transformation, selection of highly variable genes, and dimensionality reduction. Only cells that pass the cell calling step are included in these visualizations, without additional filtering. For interactive data visualization, we use plotly [16] version 6.0.0, a powerful graphing library, to generate dynamic plots directly from Python. These visualizations are exported as HTML components that maintain full interactivity in any modern web browser. We then use Beautiful Soup [17] to parse HTML templates, perform DOM manipulation, and embed the necessary components into predefined templates.

## Results

To demonstrate its utility, we applied `QCatch` to a publicly-available dataset comprising 1k human peripheral blood mononuclear cells (PBMCs) from 10X Genomics. An overview of the resulting quality control report is shown in Figure 1, and the full interactive version can be explored on the QCatch documentation site: https://combine-lab.github.io/QCatch/. In the following sections, we describe the procedure of assessing the quality of a dataset using a `QCatch` report.

**Fig. 1.**
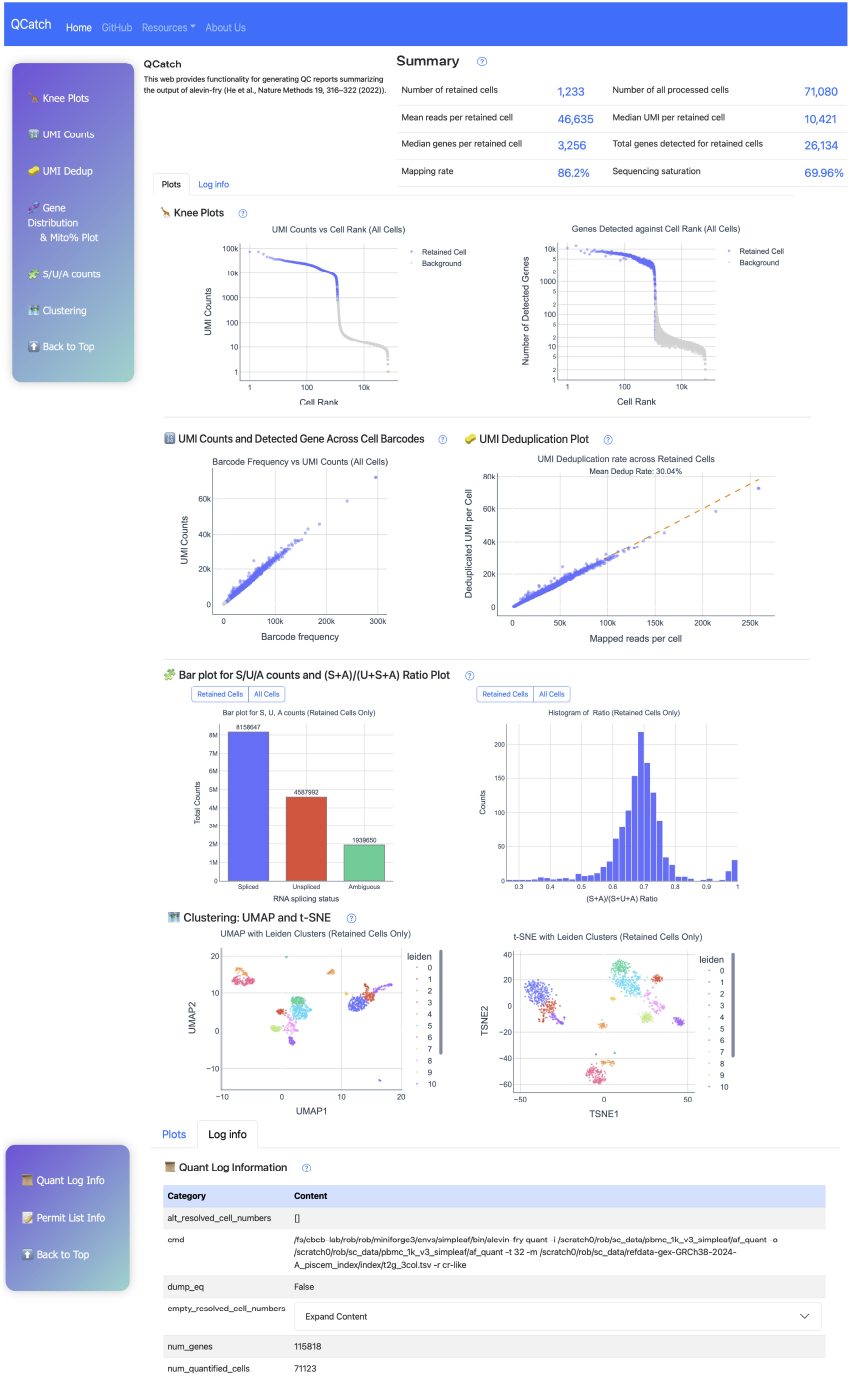
Preview of the QCatch report illustrating key components and typical content. The figure includes selectively curated and slightly reformatted sections from the original report.

### Summary statistics

The “Summary” section at the top highlights key QC metrics commonly used in single-cell data, offering a quick overview of overall data quality. The number of retained cells – estimated by QCatch’s customized cell calling algorithm – should be close to the number of cells originally loaded onto the kit. For high-quality 10X Chromium 3’ data, this typically ranges from 500 to 20,000 cells. A much higher number may indicate a failure in the cell-calling algorithm, often due to the presence of a large fraction damaged or dead cells in the sample, which is likely indicative of issues with sample quality or processing.

The number of processed cells refers to the total number of corrected cell barcodes detected in the experiment. This number is typically around 150, 000 for a 10X Chromium 3’ experiment, independent of the number of cells loaded onto the kit. A low number indicates an incomplete dataset, usually caused by processing only a subset of the FASTQ files of an experiment.

The mean reads, median UMIs, and median genes detected per retained cell reflect the sparsity of the data set and the depth of sequencing from different perspectives. For high-quality 10X Chromium 3’ datasets, we generally expect at least 20,000 reads per cell, resulting in thousands to tens of thousands of UMIs and hundreds to thousands of detected genes, depending on sequencing depth. As the sequencing depth increases, the total number of detected genes should approach the number of genes actually expressed in the sample. If these numbers are too low, deeper sequencing may be required to detect more genes. Similarly, sequencing saturation measures the number of sequenced reads per UMIs. Lower saturation implies that most reads come from distinct UMIs, suggesting potential returns with deeper sequencing. In contrast, high saturation indicates that deeper sequencing is not likely to help uncover more unique UMIs and genes.

The mapping rate represents the percentage of reads that align to the reference transcriptome. For splice-aware quantification, this includes reads mapping to both spliced transcripts and intronic regions. High-quality datasets typically have a mapping rate around 85 − 90%. A low mapping rate (e.g., below 60%) may indicate a mismatch between the sample and the reference – often due to sample contamination or using an inappropriate reference transcriptome.

These metrics collectively allow users to assess the quality of their data. For example, sample contamination or reference mismatch are suggested by low mapping rates; dead or damaged cells in the sample are suggested by an unexpectedly-high number of retained cells; Insufficient sequencing depth is suggested by low vale of mean reads per cell, median UMIs per cell, or median genes per cell, and low sequencing saturation.

### Visualizations in Plots Tab

In the Knee Plots section, we visualize both the ranked UMI counts and the number of detected genes per cell for all detected barcodes in descending order. In both plots, the x- and y-axes are shown on a log10 scale for better clarity. For high-quality data, we expect a sharp drop near the “knee” point, which corresponds to the cutoff determined by the cell-calling algorithm. This knee point separates the data into two distinct populations: one representing high-confidence barcodes with substantial read counts, and the other corresponding to potential empty droplets that may contain cell debris or ambient RNA, separated by the sharp drop. On the contrary, data with compromised quality might have a knee plot without a clear drop. In this case, in addition to high-confidence, live cells and empty droplets, there might exist other cell populations with low quality, such as dead or damaged cells. These low quality cells might be introduced in cell culturing or cell loading, or caused by the quality of the kit. When these exist, the cell calling algorithm might report a number of retained cells much higher than the number of cells loaded onto the kit.

In the UMI Counts section, we present three plots to assess cell quality and transcriptional complexity. The first two plots show barcodes (cells) ranked by total read count and plotted against two key metrics: the number of UMIs and the number of detected genes per barcode. The third plot highlights how gene detection scales with sequencing depth per cell. Barcodes with unusually high UMI counts may indicate multiplets, where a single droplet contains two or more cells. In contrast, barcodes with low read counts typically correspond to empty droplets, ambient RNA, or dead or damaged cells – suggesting low-quality barcodes that should be excluded during cell calling. Typically, high-quality data exhibit a clear positive correlation, meaning barcodes with higher read counts capture more UMIs and detect more genes, reflecting higher transcriptional complexity. While the first plot often appears linear, the last two usually show a curved trend, indicating gene detection increases rapidly with sequencing depth but eventually plateaus, reflecting the limit of gene expression in individual cells.

Further, the report compares the number of mapped reads and the number of UMIs, i.e., deduplicated reads, for each retained cell in the Barcode Collapsing Plot section. The UMI deduplication rate reflects how many unique UMIs remain after removing duplicates, compared to the total number of mapped reads.

`QCatch` introduces two novel visualizations that leverage the splicing-aware quantification capabilities of `alevin-fry` in USA mode, which categorizes UMIs into Unspliced (U), Spliced (S), and Ambiguous (A) categories. The first visualization is a bar chart that displays the total number of UMIs in each splicing category, providing a snapshot of the transcript composition across the dataset. The second is a histogram illustrating the distribution of the spliced ratio, calculated as (S + A) / (S + U + A), across all cells. This ratio represents the proportion of spliced transcripts, where ambiguous counts are typically grouped with spliced counts based on the conventional assumption that ambiguous reads, in expectation, are likely originate from spliced transcripts. If the loaded samples are nuclei, unspliced transcripts will dominate, leading to a low spliced ratio. On the contrary, when cells are loaded, most transcripts should be assigned a spliced or ambiguous splice status, since unspliced transcripts tend to stay in nuclei and to be primed less frequently.

Finally, `QCatch` provides several standard QC visualizations to support QC assessment via cell type separability and RNA composition. UMAP and tSNE are employed to provide an rough qualitative overview of the transcriptomic landscape, where each point represents a high-quality, retained cell, and spatial proximity reflects, subject to known shortcomings of these common dimensionality reduction approaches, their transcriptomic similarity – grouped cells often indicating shared cell types or states. If the expected cell types in an experiment are heterogeneous or well-differentiated, we should observe well-separated clusters in the embeddings. The presence of bridges or connecting dots between clusters may indicate doublets or multiplets, usually caused by cells containing mixed signals from more than one true cell. In such cases, detecting and removing doublets is essential to maintain data quality.

Moreover, `QCatch` generates a mitochondrial content plot showing the proportion of mitochondrial gene expression per cell. A standardized threshold of 10% mitochondrial gene expression is commonly recommended for quality control in human samples—since cells with high mitochondrial content are often damaged or dying; however, QCatch does not apply this threshold for automatic cell filtering. Instead, mitochondrial content is computed and stored in the output H5AD file as metadata, allowing users to assess data quality and make informed, dataset-specific decisions.

### Essential tables in the Log Info Tab

In addition to the interactive visualizations, the “Log info” tab presents essential information in organized tables, including the command-line arguments and parameter settings used during the run — enhancing transparency, reproducibility, and ease of reference.

## Discussion

`QCatch` provides a robust, automated solution for quality control tailored specifically to the `alevin-fry` and `simpleaf` frameworks. It streamlines the evaluation of single-cell sequencing data quality by generating comprehensive HTML reports that highlight essential metrics —including UMI distributions, cell-calling results, sequencing saturation, and RNA splicing status. By consolidating complex outputs into visually rich and well-structured summaries, `QCatch` enables researchers to efficiently assess data quality and make informed decisions for downstream analysis. Looking ahead, `QCatch` is well-positioned to evolve alongside the rapidly expanding landscape of single-cell sequencing technologies. Its modular architecture and compatibility with the scverse ecosystem facilitate seamless integration into diverse analytical workflows. Future developments will focus on extending `QCatch` ‘s functionality by introducing more modality-specific features for spatial transcriptomics, scATAC-seq, and multi-modal datasets.

In the future, we anticipate that core components of the `QCatch` ‘s QC reporting framework can be readily adapted for use in other open-source scRNA-seq quantification tools. This expansion will further enhance the availability of community-driven alternatives to the CellRanger QC reporting.

## Acknowledgment

We thank Aaron Lun from Genentech Inc. and Joshua Shapiro from Alex’s Lemonade Stand Foundation for Childhood Cancer for their insightful and valuable discussions. We also thank Yuewen Zheng from Altos Labs, Inc. for actively evaluating and testing QCatch functionalities throughout the development stage of QCatch.

This work is supported by the NIH under grant award numbers R01HG009937 to R.P. Also, this project has been made possible in part by grants DAF2024-342821, DAF2022-252586 from the Chan Zuckerberg Initiative DAF, an advised fund of Silicon Valley Community Foundation.

## Data availability

The source code and documentation of `QCatch` are available on GitHub at https://github.com/COMBINE-lab/QCatch (DOI: 10.5281 / zenodo.18134441). It can be installed via Bioconda (https://anaconda.org/bioconda/qcatch) or PyPI (https://pypi.org/project/qcatch/).

