## Supplemental File for "QCatch: A framework for quality control assessment and analysis of single-cell sequencing data"

May 31, 2025

### 1 Supplementary Figures

**Supplementary Figure 1.** Overview of the complete QCatch report layout and key elements. This figure corresponds to the full version of Figure 1 in the main manuscript, and is split across two pages for readability.

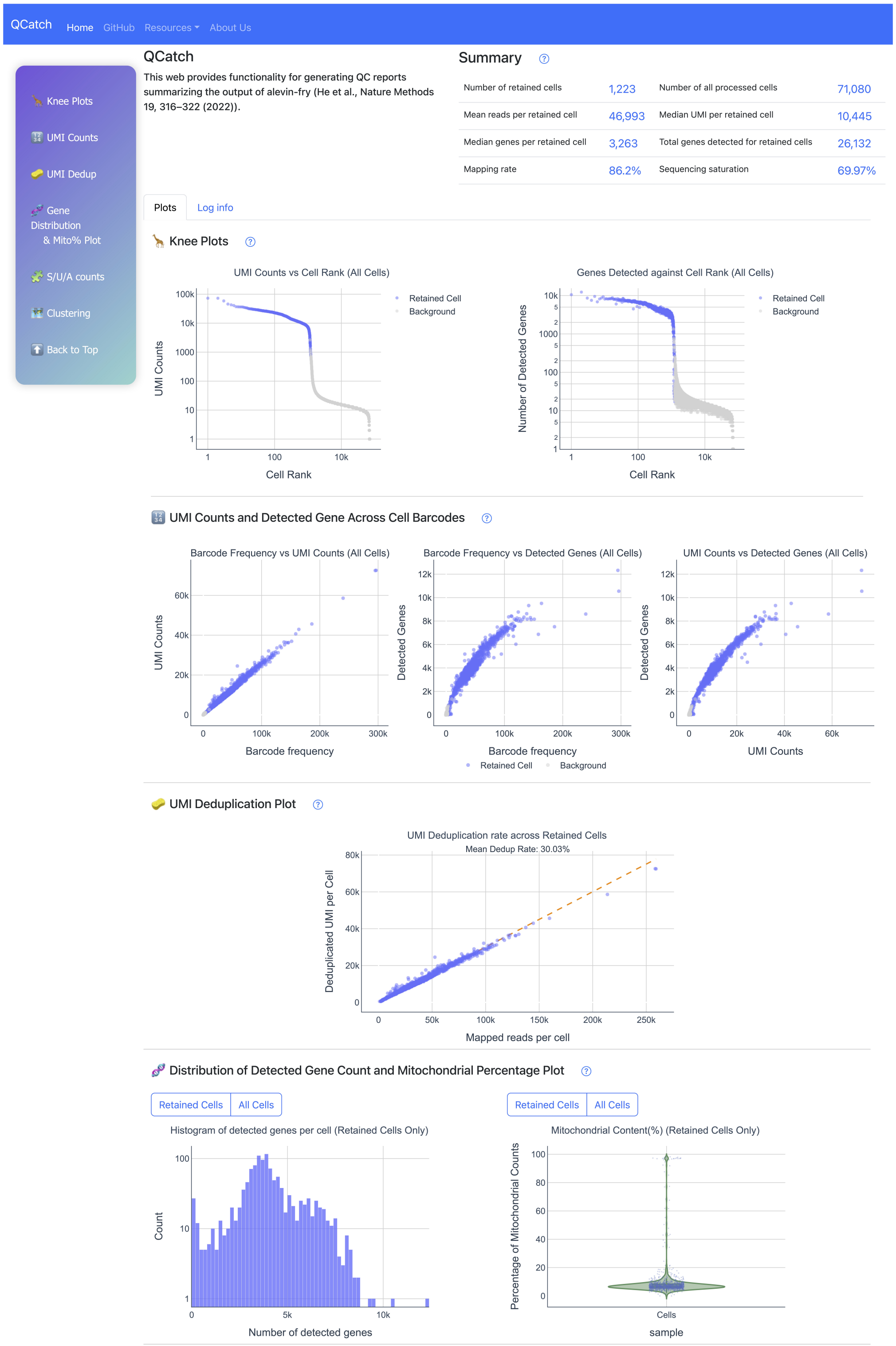

#### 💸 Bar plot for S/U/A counts and (S+A)/(U+S+A) Ratio Plot

Retained Cells | All Cells

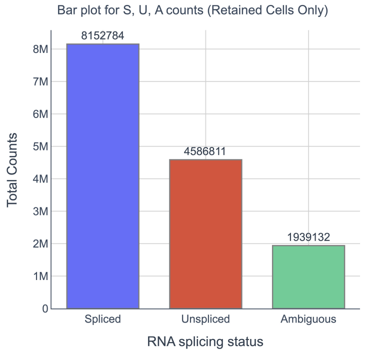


Retained Cells | All Cells

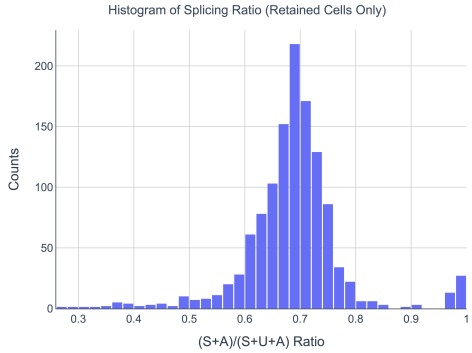


#### 🗺️ Clustering: UMAP and t-SNE

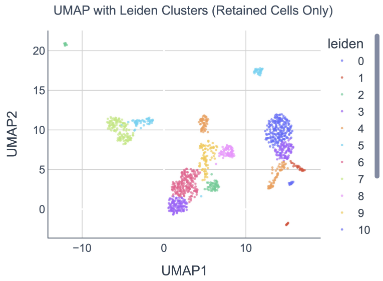


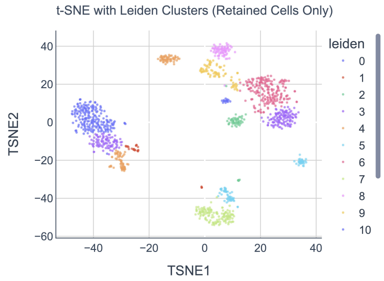

**Supplementary Figure 2.** Complete view of the "Log info" section in the QCatch report, summarizing all metadata extracted during the quantification process. This figure complements Supplementary Figure 1 by showing the structured log tables in full detail.

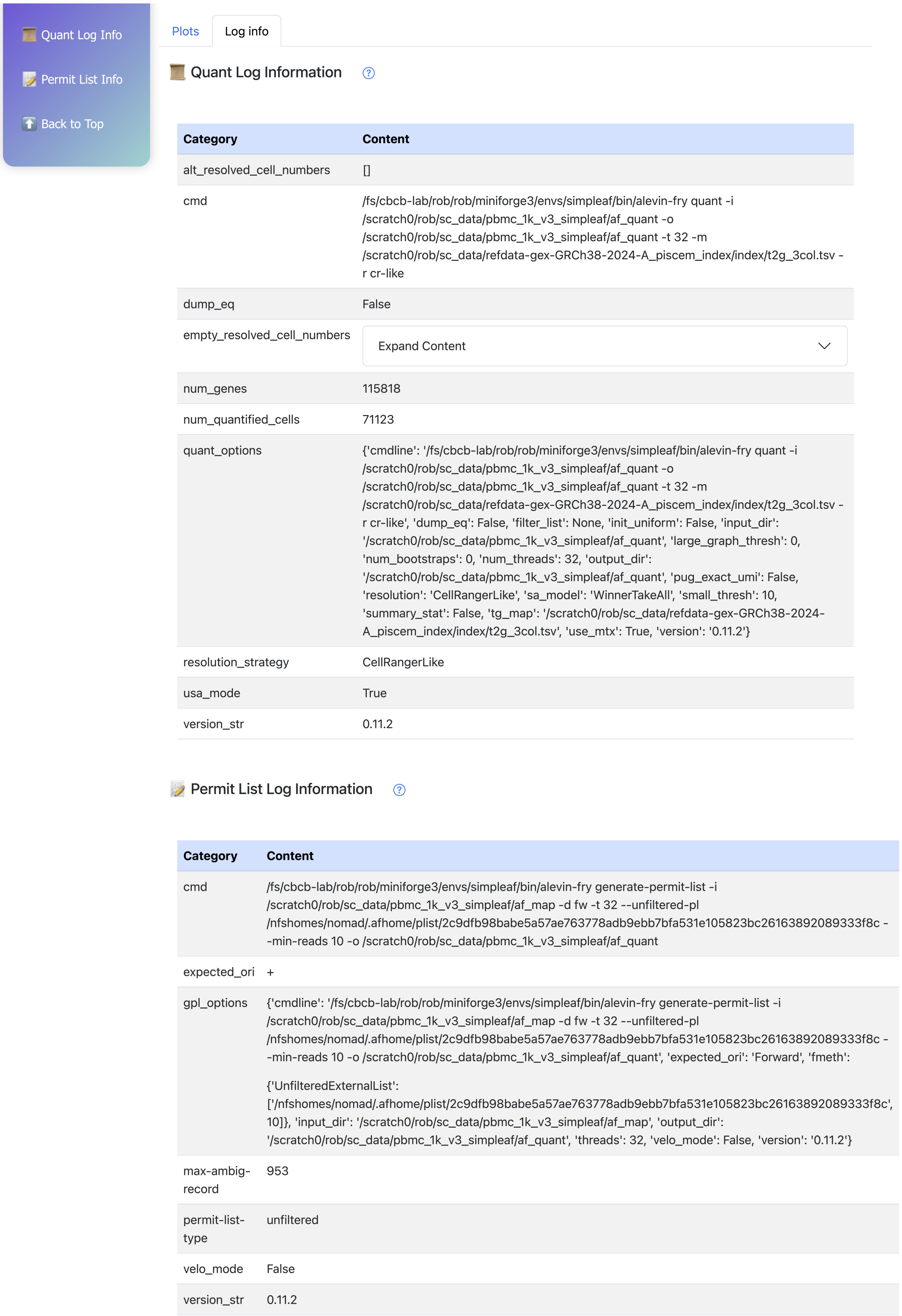



Quant Log Information

| Category | Content |
|---|---|
| alt_resolved_cell_numbers | [] |
| cmd | /fs/cbcb-lab/rob/rob/miniforge3/envs/simpleaf/bin/alevin-fry quant -i /scratch0/rob/sc_data/pbmc_1k_v3_simpleaf/af_quant -o /scratch0/rob/sc_data/pbmc_1k_v3_simpleaf/af_quant -t 32 -m /scratch0/rob/sc_data/refdata-gex-GRCh38-2024-A_piscem_index/index/t2g_3col.tsv -r cr-like |
| dump_eq | False |
| empty_resolved_cell_numbers | Expand Content |
| num_genes | 115818 |
| num_quantified_cells | 71123 |
| quant_options | {'cmdline': '/fs/cbcb-lab/rob/rob/miniforge3/envs/simpleaf/bin/alevin-fry quant -i /scratch0/rob/sc_data/pbmc_1k_v3_simpleaf/af_quant -o /scratch0/rob/sc_data/pbmc_1k_v3_simpleaf/af_quant -t 32 -m /scratch0/rob/sc_data/refdata-gex-GRCh38-2024-A_piscem_index/index/t2g_3col.tsv -r cr-like', 'dump_eq': False, 'filter_list': None, 'init_uniform': False, 'input_dir': '/scratch0/rob/sc_data/pbmc_1k_v3_simpleaf/af_quant', 'large_graph_thresh': 0, 'num_bootstraps': 0, 'num_threads': 32, 'output_dir': '/scratch0/rob/sc_data/pbmc_1k_v3_simpleaf/af_quant', 'pug_exact_umi': False, 'resolution': 'CellRangerLike', 'sa_model': 'WinnerTakeAll', 'small_thresh': 10, 'summary_stat': False, 'tg_map': '/scratch0/rob/sc_data/refdata-gex-GRCh38-2024-A_piscem_index/index/t2g_3col.tsv', 'use_mtx': True, 'version': '0.11.2'} |
| resolution_strategy | CellRangerLike |
| usa_mode | True |
| version_str | 0.11.2 |

Permit List Log Information

| Category | Content |
|---|---|
| cmd | /fs/cbcb-lab/rob/rob/miniforge3/envs/simpleaf/bin/alevin-fry generate-permit-list -i /scratch0/rob/sc_data/pbmc_1k_v3_simpleaf/af_map -d fw -t 32 --unfiltered-pl /nfshomes/nomad/.afhome/plist/2c9dfb98babe5a57ae763778adb9ebb7bfa531e105823bc26163892089333f8c --min-reads 10 -o /scratch0/rob/sc_data/pbmc_1k_v3_simpleaf/af_quant |
| expected_ori | + |
| gpl_options | {'cmdline': '/fs/cbcb-lab/rob/rob/miniforge3/envs/simpleaf/bin/alevin-fry generate-permit-list -i /scratch0/rob/sc_data/pbmc_1k_v3_simpleaf/af_map -d fw -t 32 --unfiltered-pl /nfshomes/nomad/.afhome/plist/2c9dfb98babe5a57ae763778adb9ebb7bfa531e105823bc26163892089333f8c --min-reads 10 -o /scratch0/rob/sc_data/pbmc_1k_v3_simpleaf/af_quant', 'expected_ori': 'Forward', 'fmeth': {'UnfilteredExternalList': ['/nfshomes/nomad/.afhome/plist/2c9dfb98babe5a57ae763778adb9ebb7bfa531e105823bc26163892089333f8c', 10]}, 'input_dir': '/scratch0/rob/sc_data/pbmc_1k_v3_simpleaf/af_map', 'output_dir': '/scratch0/rob/sc_data/pbmc_1k_v3_simpleaf/af_quant', 'threads': 32, 'velo_mode': False, 'version': '0.11.2'} |
| max-ambig-record | 953 |
| permit-list-type | unfiltered |
| velo_mode | False |
| version_str | 0.11.2 |

### 2 Supplementary Documentations

#### 2.1 Explanation of Metadata Column Names in `anndata.obs`

- **CB:** Cell Barcode
- **Corrected Reads:** The total number of corrected reads, including both mapped and unmapped reads.
- **Mapped Reads:** The number of reads that were successfully mapped to the reference.
- **Deduplicated Reads:** The number of deduplicated reads, processed by UMI deduplication.
- **Mapping Rate:** The ratio of mapped reads to corrected reads.
- **Deduplication Rate:** The ratio of deduplicated reads to mapped reads.
- **Mean by Max:** Calculated as mean_expr / max_umi, where:
  **mean_expr** is the total UMI count divided by the number of genes with non-zero expression levels.
  **max_umi** is the maximum expression level across all genes.
- **Number of Genes Expressed:** The number of genes expressed in this cell.
- **Number of Genes Above Mean:** The number of genes with expression levels above the mean_expr, where mean_expr is defined as the total UMI count divided by the number of genes with non-zero expression levels.

#### 2.2 Summary Metrics

- **Number of retained cells:** The number of valid and high quality cells that passed the cell calling step. This includes cells identified during the initial filtering and additional cells identified by the EmptyDrops step, whose expression profiles are significantly distinct from the ambient background.
- **Number of all processed cells:** The total number of cell barcodes observed in the processed sample. Cells with zero reads have been excluded.
- **Mean reads per retained cell:** The total number of reads assigned to the retained cells, including the mapped and unmapped reads, divided by the number of retained cells.
- **Median UMI per retained cell:** The median number of deduplicated reads (UMIs) per retained cell.
- **Median genes per retained cell:** The median number of genes detected per retained cell.
- **Total genes detected for retained cells:** The total number of unique genes detected across all retained cells.
- **Mapping rate:** Fraction of reads that mapped to the reference, calculated as mapped reads / total processed reads.
- **Sequencing saturation:** Sequencing saturation measures the proportion of reads coming from already-seen UMIs, calculated as 1 - (deduplicated reads / total reads). High saturation suggests limited gain from additional sequencing, while low saturation indicates that further sequencing could reveal more unique molecules (UMIs).

##### 2.3 Plots

This section describes how each plot in the QCatch report is generated from the processed single-cell quantification data.

**Knee plots**
We order the UMI count (deduplicated reads) for each cell and sort in descending order to get the cell rank, and use the scatter plot to display the UMI count against the cell rank. We also use the scatter plot to display the number of detected genes against the cell rank.

**UMI Counts and Detected Gene Across Cell Barcodes**
The barcode frequency is calculated as the number of reads associated with each cell barcode. The first two plots show barcode frequency against two key metrics: the number of UMIs and the number of detected genes per barcode. The third plot illustrates how the number of detected genes increases with UMI count per cell.

**UMI Deduplication plot**
The scatter plot compares the number of mapped reads and number of UMIs for each retained cell. Each point represents a cell, with the x-axis showing the mapped reads count and the y-axis showing the deduplicated UMIs count. The reference line indicates the mean deduplication rate across all cells.
**UMI Deduplication:** UMI deduplication is the process of identifying and removing duplicate reads that arise from PCR amplification of the same original molecule.
**Dedup Rate:** The UMI count divided by the number of mapped reads for each cell.

**Distribution of Detected Gene Count and Mitochondrial Percentage Plot**
The plot depicts the distribution of detected gene counts per cell. The violin plot shows the distribution of mitochondrial gene expression percentages.
**Note:** The “All Cells” plot does not display every processed cell. To improve visualization and reduce clutter from very low-quality cells, we excluded cells with fewer than 20 detected genes—these are typically considered nearly empty. In contrast, the “Retained Cells” plot includes all retained cells, without applying this gene count filter.

**Bar Plot for S/U/A Counts and Splicing Ratio Distribution**
When using “USA mode” in alevin-fry, spliced (S), unspliced (U), and ambiguous (A) read counts are generated separately for each gene in each cell.
In the bar plot, we first sum the spliced, unspliced, and ambiguous counts across all genes and all cells. The plot then displays the total number of reads in each splicing category: Spliced (S), Unspliced (U), and Ambiguous (A).
In the histogram, we calculate the splicing ratio for each cell as (S + A) / (S + U + A), where the counts are summed across all genes. The histogram shows the distribution of these per-cell splicing ratios.

**Clustering: UMAP and t-SNE**
These plots represent low-dimensional projections of high-dimensional gene expression profiles. Each point corresponds to a single cell. Cells that are positioned close to one another in the plot are inferred to have similar transcriptomic signatures, suggesting similarity in cell type or state.

**Note:** Only retained cells are included in these visualizations, and no additional filtering was applied beyond the cell retention step. Standard preprocessing was performed using Scanpy, including normalization, log transformation, highly variable gene selection, and dimensionality reduction.
